# Stromal cell-contact dependent PI3K and APRIL induced NF-κB signaling complement each other to prevent mitochondrial- and endoplasmic reticulum stress induced cell death of bone marrow plasma cells

**DOI:** 10.1101/849638

**Authors:** Rebecca Cornelis, Stefanie Hahne, Adriano Taddeo, Georg Petkau, Darya Malko, Pawel Durek, Manja Thiem, Lukas Heiberger, Elodie Mohr, Cora Klaeden, Koji Tokoyoda, Francesco Siracusa, Bimba Franziska Hoyer, Falk Hiepe, Mir-Farzin Mashreghi, Fritz Melchers, Hyun-Dong Chang, Andreas Radbruch

## Abstract

Persistence of long-lived, memory plasma cells in the bone marrow depends on survival factors available in the bone marrow, provided in niches organized by stromal cells. Here we describe that *ex vivo* we can prevent apoptosis of bone marrow plasma cells by supplying direct cell contact with stromal cells and the soluble cytokine APRIL. Integrin-mediated contact of bone marrow plasma cells with stromal cells activates the PI3K signaling pathway, leading to critical inactivation of FoxO1/3 and preventing the activation of mitochondrial stress-associated effector caspases 3 and 7. Likely, inhibition of PI3K signaling *in vivo* ablates bone marrow plasma cells. APRIL signaling, via the NF-κB pathway, blocks activation of the endoplasmic reticulum stress-associated initiator caspase 12. Thus, stromal cell-contact induced PI3K and APRIL-induced NF-κB signaling provide necessary and complementary signals to maintain bone marrow memory plasma cells.

## Introduction

Plasma cells can persist for long time periods in the bone marrow. However, plasma cells are not intrinsically long-lived (Makela & Nossal, 1962; Schooley, 1961) and die quickly when isolated and cultured *in vitro*, suggesting that their persistence in the bone marrow depends on survival factors provided in the bone marrow (Cassese et al., 2003). In the bone marrow, plasma cells are located individually in direct contact to mesenchymal stromal cells, and it has been postulated that these stromal cells organize a survival niche for the plasma cell (Manz & Radbruch, 2002; Radbruch et al., 2006; Tokoyoda, Egawa, Sugiyama, Choi, & Nagasawa, 2004; Zehentmeier et al., 2014). How the survival is mediated at a molecular level has remained unclear. Essential for plasma cell survival is signaling through the B cell maturation antigen (BCMA, CD269) receptor of the plasma cell (O’Connor et al., 2004a) induced by its two ligands a proliferation-inducing ligand (APRIL, CD256) or B cell activating factor (BAFF, BLys, CD257) (Benson et al., 2008), and the anti-apoptotic protein myeloid cell leukemia 1 (MCL-1) (Peperzak et al., 2013). There is evidence that stromal cells might contribute directly to plasma cell survival in the bone marrow. Antibodies against the adhesion molecules integrin αLβ2 (lymphocyte function-associated antigen, LFA-1, CD11a/CD18) and integrin α4β1 (very late antigen, VLA-4, CD49d/CD29), expressed by the plasma cells, ablate plasma cells from the bone marrow (DiLillo et al., 2008). Ligands for both of these integrins, vascular cell adhesion molecule 1 (VCAM, CD106) and intercellular adhesion molecule 1 (ICAM, CD54), are expressed by bone marrow stromal cells, suggesting that integrin-mediated binding of plasma cells to stromal cells might directly, via the focal adhesion kinase/Phosphatidylinositol-4,5-bisphosphate 3-kinase (PI3K) pathway (Giancotti, 1999; Parsons, Martin, Slack, Taylor, & Weed, 2000) promote plasma cell persistence in the bone marrow. In line with this, CD37-deficient antibody secreting cells, which show impaired clustering of VLA-4 have diminished PI3K signaling and impaired survival (Spriel et al., 2012). Whether or not BCMA signaling and contact to stromal cells are sufficient to maintain survival of memory plasma cells, and how they prevent death of the plasma cells, has not been elucidated so far.

Here we demonstrate both *in vivo* and *ex vivo*, that bone marrow plasma cell survival is dependent on PI3K signaling. PI3K signaling is induced in plasma cells by direct contact to stromal cells and leads to inactivation of Forkhead-Box-Protein O1/3 (FoxO1/3), which is essential for the survival of the plasma cells. Plasma cell survival also dependents on nuclear factor ‘kappa-light-chain-enhancer’ of activated B-cells (NF-κB) signaling, that is induced by APRIL. Pan-caspase inhibition can substitute for both signaling pathways and rescues plasma cell survival *in vitro*. Interestingly, stromal cell contact alone but not APRIL prevents activation of the mitochondrial stress-associated caspases 3 and 7, whereas APRIL prevents activation of the endoplasmic reticulum (ER)-associated caspase 12.

## Results

### Contact to stromal cells and APRIL-induced signaling pathways prevent caspase-mediated cell death of plasma cells

Memory plasma cells were isolated from the bone marrow of chicken gamma globulin (CGG) immunized C57BL/6J mice more than 30 days after the last immunization, by magnetic depletion of CD49b^+^ and B220^+^ cells, followed by magnetic enrichment of CD138^+^ cells. Using this protocol, purities of more than 90% and recovery rates of about 35% of viable bone marrow plasma cells were achieved. Isolated plasma cells expressed the plasma cell transcription factor BLIMP-1 and were Ki-67 negative, the latter indicating that the cells were resting in terms of proliferation (suppl. Figure 1A,B). The cells were cultured *in vitro* with or without murine stromal cell line ST2 at an initial ratio of 1:1 in the presence or absence APRIL. On days 1, 3 and 6 of the culture, viable plasma cells (CD138^++^/DAPI^-^) were enumerated and analyzed by flow cytometry. All cultures were performed under physiological oxygen levels of 4.2% O_2_ to mimic the bone marrow environment (Spencer et al., 2014). Plasma cells rapidly died within days, when isolated from the bone marrow and cultured in RPMI 1640/FCS (median viability: d1: 43.27%/ d3: 7.095%/ d6: 0%). However, plasma cell survival was significantly improved, when the cells were co-cultured with ST2 stromal cells and in the presence of the cytokine APRIL (median viability: d1: 83.14%/ d3: 72.19%/ d6: 51.20%). Co-culture of plasma cells with ST2 cells alone (median viability: d1: 67.47%/ d3: 25.42% / d6: 19.07%) or with APRIL alone (median viability: d1: 55.24% / d3: 43.15%/ d6: 23.27%) were not sufficient to maintain plasma cells alive (Figure 1A). The expression of CD138 and BLIMP-1 on the plasma cells was not altered during the 6 days of *in vitro* culture with ST2 stromal cells and APRIL, and antibody secretion was maintained (suppl. Figure 1C,D). To confirm that the identity of plasma cells was maintained for 3 days in co-culture with ST2 stromal cells and APRIL, we compared their global transcriptomes to those of *ex vivo* isolated plasma cells (suppl. Figure 1E, F). Of 10.000 genes expressed at statisticaly significant levels, only 41 genes showed a significant difference (p < 0.01) in expression before and after culture (suppl. Figure 2). The transcription factor AP-1 (JunB, Jun, Fos) was highly expressed in plasma cells isolated from bone marrow, but not or only marginally in plasma cells cultivated for 3 days. Expression of AP-1 and other stress inducible genes (12, group A) may reflect stress induced by the tedious isolation procedure of plasma cells from the bone marrow, as compared to their isolation from cell culture. A number of hypoxia-related and metabolic genes (15 genes, group B) were upregulated in 3 days cultivated plasma cells, as compared to plasma cells directly isolated from the bone marrow. This may reflect the extended time the plasma cells had spent in normoxic conditions during the isolation period. Finally, other genes (12 genes, group C), most prominently CXCL12, not expressed in plasma cells isolated direcly from the bone marrow, but in those isolated from cell culture on day 3, may indicate contaminating stromal cells which were abundant in cell culture due to the ST2 cell line. Expression of the genes *CD138, Foxo1* and *3, Prdm1, Irf4, Noxa, Bcl2l11, Bcl2* and *Mcl1* was not significantly different (suppl. Figure 1 G).

**Figure 1.**
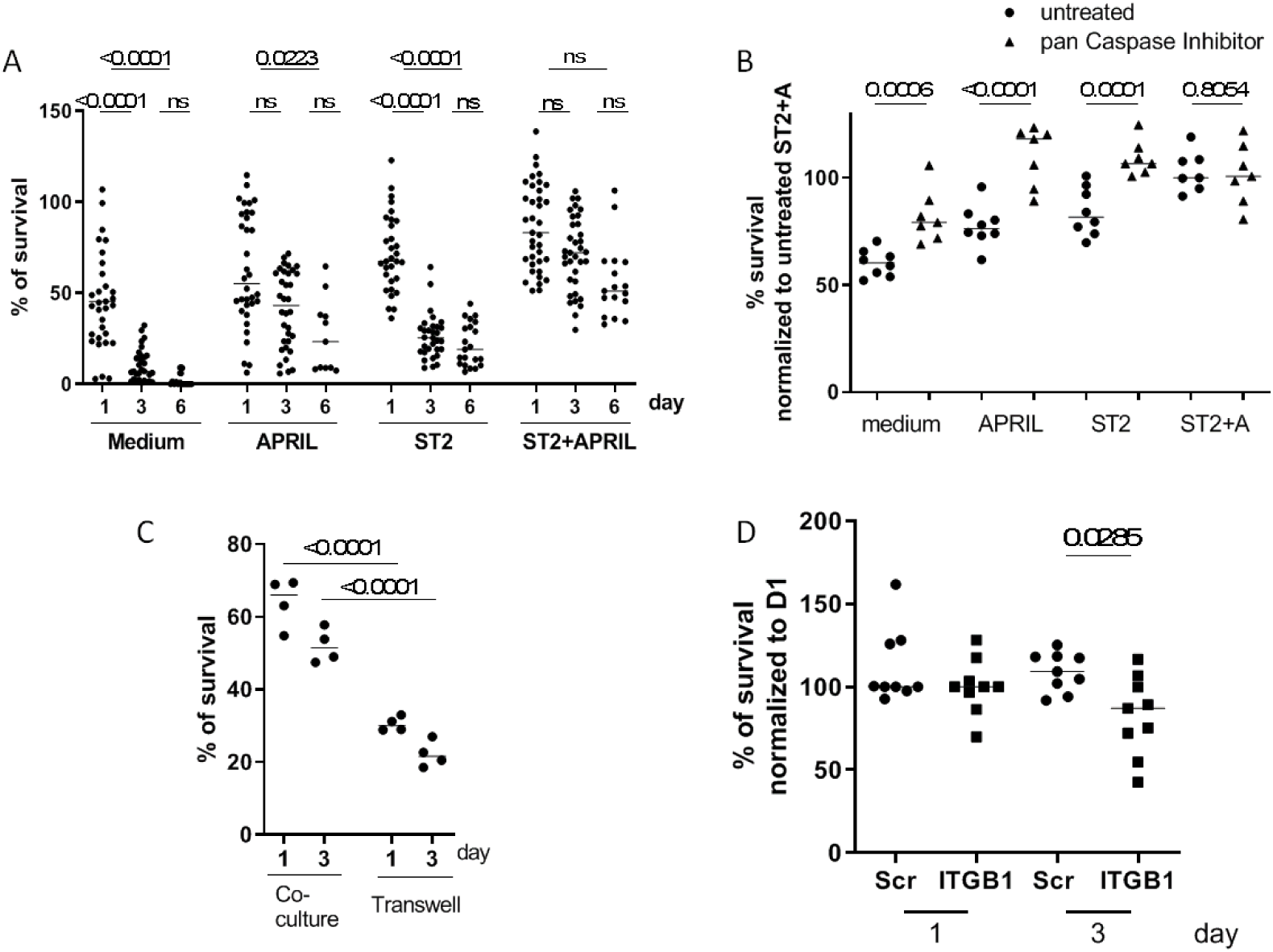
Survival of bone marrow memory plasma cells is dependent on direct cell contact with stromal cells and the presence of APRIL. (A) Survival of primary plasma cells isolated from immunized C57BL/6J mice cultured with or without ST2 stromal cell lines and with or without APRIL for up to 6 days at 4.2% O_2_. Viable plasma cells (CD138++/DAPI-) were enumerated by flow cytometry and data are represented as median (pooled from a minimum of 5 independent experiments with a minimum of n=14 for each group). (B) Isolated plasma cells were treated with or without pan-caspase inhibitor when cultured with or without ST2 stromal cell lines and with or without APRIL. Viable plasma cells were counted on day 1 of culture (pooled from two independent experiments with a minimum of n=7 for each group). (C) Survival of plasma cells in the presence of APRIL on day 1 and day 3, when cultured in transwell or in direct contact to stromal cells (pooled from two independent experiments with n=4 for each group). (D) Survival of plasma cells on day 1 and day 3 treated with specific siRNA directed against ITGB1 and scrambled controls (pooled from three independent experiments with n=9 for each group).

The survival of *ex vivo* isolated bone marrow plasma cells cultured with APRIL or ST2 stromal cells alone was rescued by pan-caspase inhibitors (Fig. 1B), suggesting that co-culture of plasma cells with ST2 stromal cells and APRIL prevents caspase-mediated cell death.

Interaction between stromal cells and plasma cells has been suggested to be mediated by direct cell contact (Manz & Radbruch, 2002; Radbruch et al., 2006; Tokoyoda et al., 2004; Zehentmeier et al., 2014) via VCAM1/VLA4 and ICAM/LFA1 interaction (DiLillo et al., 2008). When culturing plasma cells and ST2 stromal cells in a transwell assay, survival of plasma cells was significantly decreased (median viability: d1: 66% / d3: 51% and d1: 30%/ d3: 22%, respectively) as compared to co-culturing conditions (Fig. 1C). siRNA-mediated knock-down of integrin β1 (CD29/ITGB1), a subunit of the VLA4 (Integrin a4ß1) heterodimer, significantly reduced the survival of plasma cells in co-culture with ST2 stromal cells and APRIL *ex vivo* (Fig. 1D), indicating that direct cell contact is required for survival and that contact-mediated survival is in part mediated by integrin β1 (median viability: scr d1: 100%, d3: 109% / ITGB1 d1: 100%, d3: 87%).

### Inhibition of PI3K signaling results in plasma cell death ex vivo

Preincubation of ex vivo isolated bone marrow plasma cells with the irreversible PI3K inhibitor Wortmannin resulted in a dose-dependent decrease in survival of the plasma cells, when co-cultured with ST2 stromal cells in the presence of APRIL (viability: ST2+A: d1 101%, d3 76% /+Wortmannin 0.6 μM: d1 100% d3 66% / +Wortmannin 3 μM: d1 102%, d3 66% / +Wortmannin 15 μM: d1 73%, d3 37% / +Wortmannin 77 μM: d1 68%, d3 20%) (Figure 2A). Using the pan-PI3K inhibitor, LY294002, plasma cell survival was decreased to a similar degree (viability: ST2+A, +LY29400 10 μM: d1 62%, d3 68% / +LY29400 20 μM: d1 44%, d3 37% / +LY29400 40 μM: d1 24%, d3 5.7%) (Fig. 2B). More specific inhibition of any of the four known PI3K subunits α, β, γ or δ did not impact the survival of plasma cells (suppl. Fig. 3). Only when any three subunits were simultaneously inhibited, plasma cell survival was reduced to a similar degree as it was observed in the presence of Wortmannin (Fig. 2C). Apparently, plasma cells have no particular requirement regarding the subunit composition of their PI3K. Inhibition of the NF-κB pathway downstream of BCMA, the receptor for APRIL, with the pan-NF-κB inhibitor IKK16 (Thein, Pham, Bayer, Tao-Cheng, & Dosemeci, 2008; Waelchli et al., 2006) also resulted in a dose-dependent death of the plasma cells (suppl. Fig. 4), demonstrating that both signaling pathways downstream of stromal cell contacts and BCMA are non-redundant and essential for plasma cell survival.

**Figure 2.**
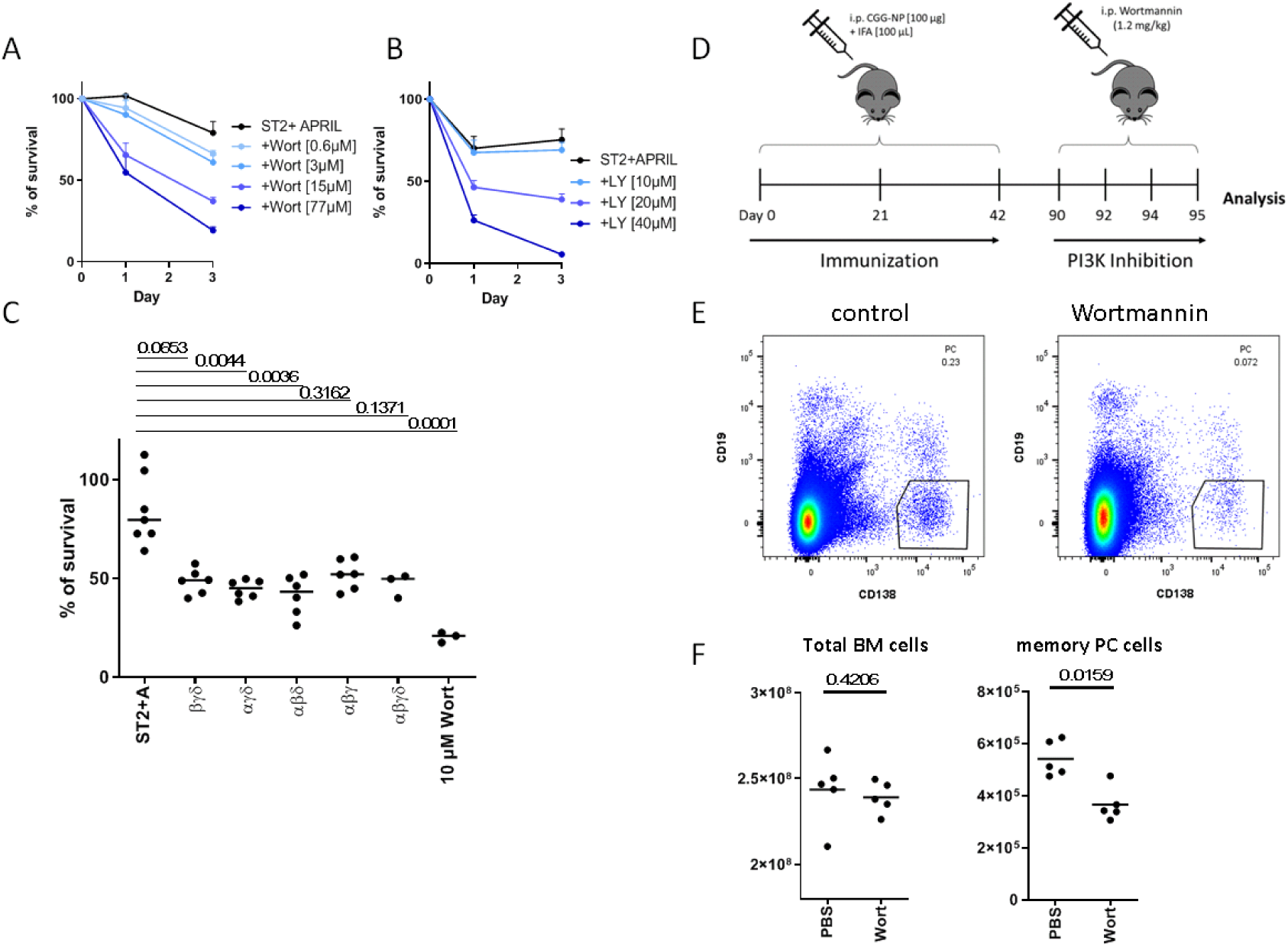
Stromal cell contact-induced PI3K signaling is essential for survival of memory bone marrow plasma cells *ex vivo* and *in vivo*. Survival of plasma cells (A) pre-incubated with different concentrations of the irreversible PI3K-inhibitor Wortmannin or (B) directly treated during culture with the inhibitor LY294002 or (C) with combinations of three subunit-specific PI3K-inhibitors determined by enumeration of viable CD138^++^/DAPI^-^ plasma cells by flow cytometry (pooled from two independent experiments with n=6 for each group). (D) Experimental design: C57BL/6J were primed and boosted twice (days 21 and 42) with CGG-NP/IFA and treated with the PI3K-inhibitor Wortmannin on days 90, 92, and 94. On day 95, the mice were analyzed. (E) Representative plot of B220^-^/CD138^++^/CD19^-^ plasma cells in the bone marrow from control and Wortmannin-treated mice gated on DAPI^-^ viable cells. (F) Absolute cell counts of total bone marrow cells and memory plasma cells, in the bone marrow in control and Wortmannin-treated mice, data are represented as median (pooled from two independent experiments with n=5).

### Inhibition of PI3K signaling ablates resident memory plasma cells of the bone marrow

To verify the relevance of PI3K-signaling for the persistence of plasma cells *in vivo*, we treated mice with an established immune memory with 1.2 mg/kg of Wortmannin (Nobs et al., 2015), 7 weeks following the last immunization and enumerated memory plasma cells in the bone marrow on day 95 (Figure 2D, E). Counts of total bone marrow cells did not differ between mice treated with Wortmannin and control mice (2.46×10^8^ versus 2.37×10^8^) (Figure 2F). However, plasma cells were significantly reduced by about 67% (5.1×10^5^ versus 3.4×10^5^).These results show that persistence of memory plasma cells in the bone marrow, like in the *ex vivo* niche provided by ST2 stromal cells and APRIL, is conditional on continued PI3K-signaling.

### Stromal cell contact downregulates the FoxO1/3 pathway

PI3K activation leads to the downregulation of the transcription factors forkhead box protein O1 and O3 (FoxO1 and 3) (Haftmann et al., 2012; Huang et al., 2005; Plas & Thompson, 2003). Bone marrow plasma cells, when co-cultured with ST2 stromal cells significantly downregulated the expression of FoxO1 and FoxO3 independently of APRIL, already on day 1 of co-culture (FoxO1 geometric mean expression:, APRIL: 1820 ± 62, ST2: 1374 ± 76, ST2+A: 1348 ± 35/ FoxO3a geometric mean expression:, APRIL: 2446 ± 282, ST2: 1777 ± 134, ST2+A: 1960 ± 106) (Figure 3A, B). Adding APRIL alone or in combination with ST2 cells did not affect the expression of FoxO1/3 proteins. To determine whether downregulation of FoxO1/3 expression is the critical event downstream of PI3K activation, FoxO1/3 expression was knocked-down using specific siRNA. Knock-down of FoxO1/3 in plasma cells could completely restore plasma cell survival in the absence of ST2 stromal cells when plasma cells were cultured with APRIL alone (mean viability: day1, APRIL+ scr: 99% ± 13, APRIL+ FoxO: 102% ± 14, ST2+A+scr: 101% ± 33/ day3, APRIL+ scr: 52% ± 26, APRIL+ FoxO: 79% ± 18, ST2+A+scr: 89% ± 24) (Figure 3C). These results demonstrate that stromal cell contact is supporting survival of plasma cells by downregulation of FoxO1/3.

**Figure 3.**
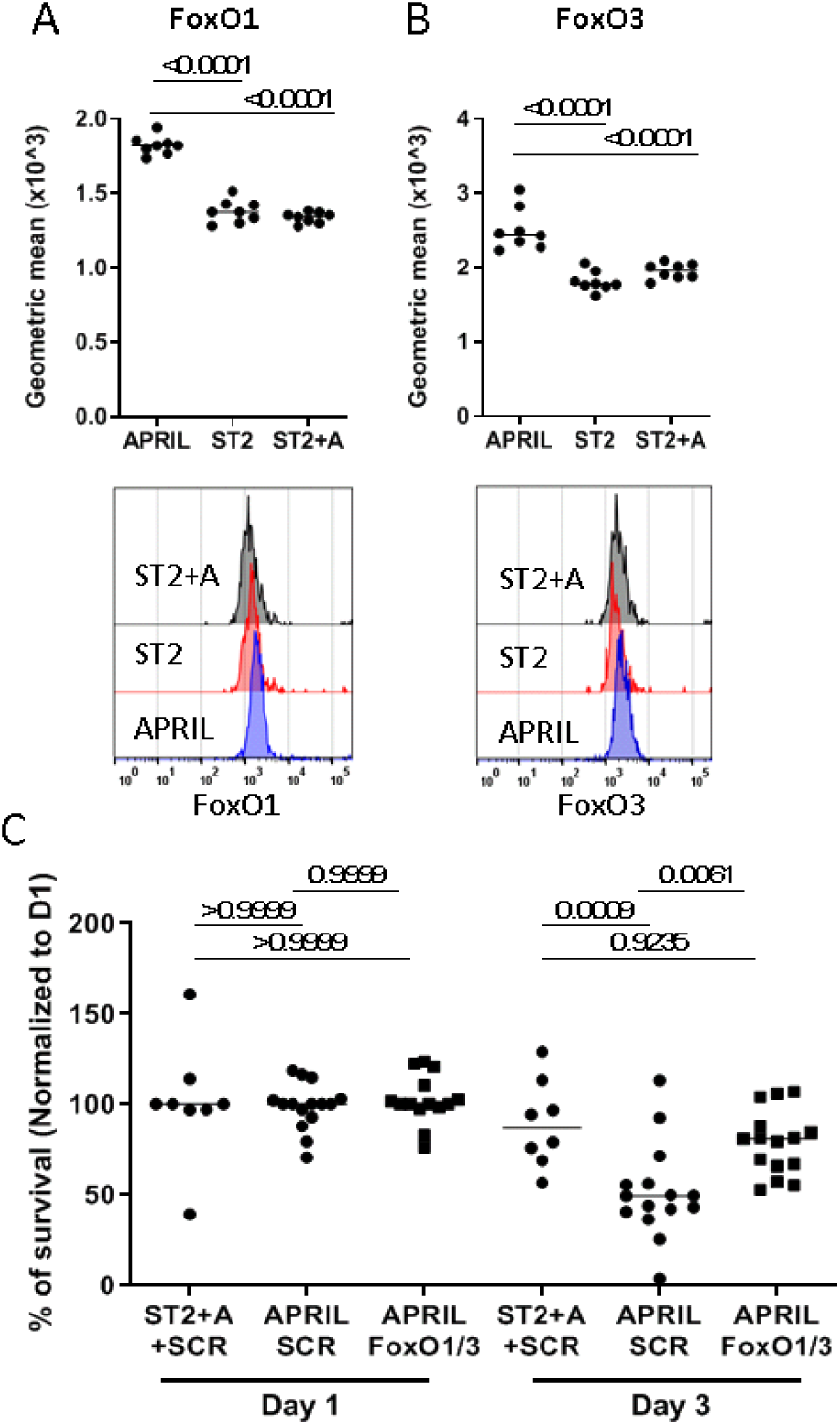
Stromal cell contact-induced PI3K signaling downregulates FoxO1/3 protein expression and is essential for survival of memory bone marrow plasma cells. FoxO1 (A) and FoxO3 (B) expression as determined by flow cytometry (geometric mean fluorescence intensity, pooled from two independent experiments with n=8 for each group). (C) Survival of plasma cells treated with siRNA specific for FoxO1/FoxO3 or scrambled control on day 1 and 3 of culture with APRIL alone or ST2 and APRIL as enumerated by flow cytometry. Data are represented as median (pooled from a minimum of three independent experiments with n=3 for each group).

### Stromal cell contact and APRIL signaling pathways address distinct caspases

As the survival of *ex vivo* isolated bone marrow plasma cells cultured with APRIL or ST2 stromal cells alone was rescued by pan-caspase inhibitors (Figure 1B), we aimed at determining which caspases are affected by stromal cell contact and APRIL, respectively. Activation of the caspases was measured on the single-cell level using fluorescent caspase-specific peptides. Co-culture of plasma cells with ST2 stromal cells led to significantly reduced levels of cleavage and activation of the effector caspases 3 and 7 (Figure 4A, B). APRIL by itself or in combination with ST2 stromal cells did not impact on activation of caspase 3 or 7. APRIL, together with ST2 stromal cells, led to a significant reduction of the activation of the ER-associated caspase 12 (Figure 4C).

**Figure 4.**
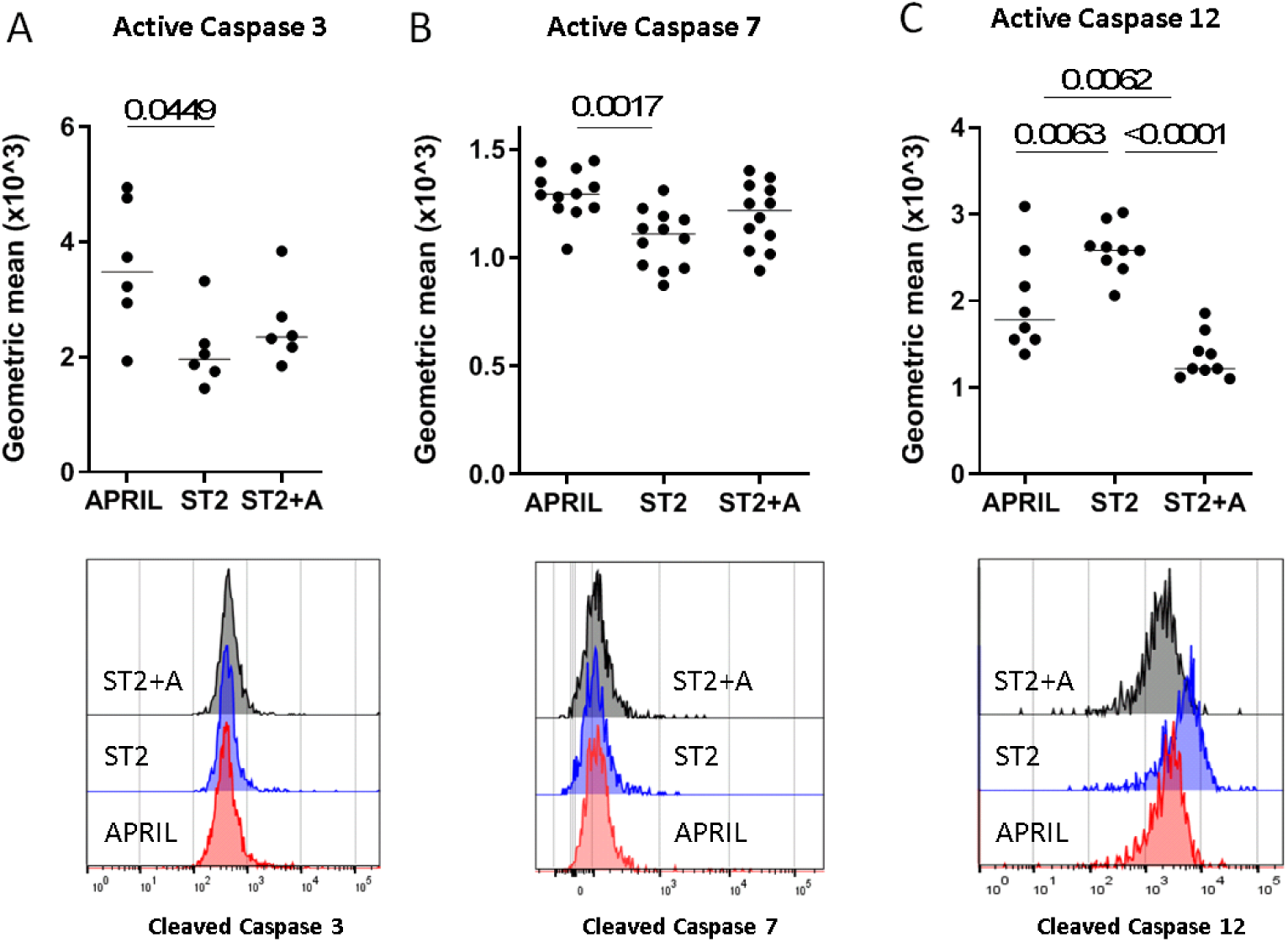
Contact with stromal cells inhibits activation of caspase 3 and 7, while APRIL inhibits activation of caspase 12. Expression of activated (A) caspase 3, (B) caspase 7 and (C) caspase 12 determined by flow cytometry, shown as geometric mean fluorescence intensity, on day 1 in plasma cells cultured under the indicated conditions (pooled from a minimum of two independent experiments with n=6-12 for each group).

### Stromal cell contact and APRIL synergize to induce the expression of IRF4

ST2 stromal cells and APRIL also synergized to upregulate the expression of IRF4 on the single cell level (Fig. 5C). IRF4 has been demonstrated previously to be indispensable for the survival of plasma cells *in vivo* (Tellier, 2016). Inhibition of either signaling pathways downstream of stromal cell contact or APRIL, i.e. PI3K using Wortmannin or NF-κB using IKK16 resulted in a decrease in IRF4 expression (Fig. 5).

**Figure 5.**
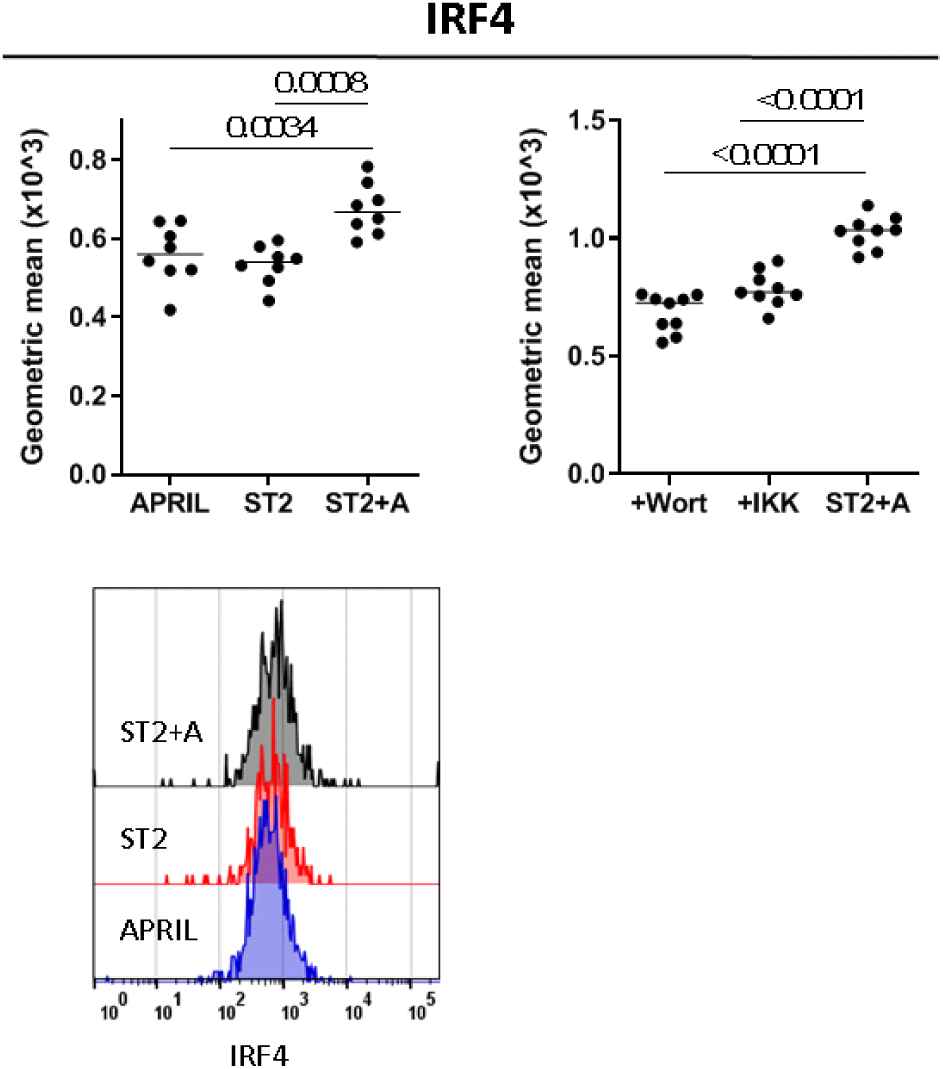
Both PI3K and NF-κB signaling synergize to upregulate expression of IRF4 in memory plasma cells. IRF4 protein expression, shown as geometric mean, in plasma cells cultured for 1 day with or without ST2 cells, with or without APRIL or pre-treated with either 10 μM Wortmannin or 2.5 μM of IKK16 (pooled from two independent experiments with n=6-8 for each group).

## Discussion

Long-lived memory plasma cells persist for a lifetime in mice (Manz, Thiel, & Radbruch, 1997; Slifka, Antia, Whitmire, & Ahmed, 1998), non-human primates (Hammarlund et al., 2017) and in humans (Landsverk et al., 2017). They form an independent population of memory cells (Chang, Tokoyoda, & Radbruch, 2018; Radbruch et al., 2006). Memory plasma cells are evenly distributed throughout the bone marrow and most if not all of them directly contact a mesenchymal stromal cell (Zehentmeier et al., 2014). It is suggestive that these stromal cells organize a survival niche, which supports longevity of the plasma cell (Manz & Radbruch, 2002). It is still enigmatic, however, how stromal cells organize the plasma cell survival niche. It has been postulated, that they attract cells secreting the cytokines APRIL and/or BAFF, both ligands for the receptor B cell maturation antigen (BCMA, CD269), signaling via the NF-κB pathway (Chu et al., 2011; Hatzoglou et al., 2000; O’Connor et al., 2004b). Blocking both cytokines ablates plasma cells from the bone marrow (Benson et al., 2008). The stromal cell may also contribute directly to the survival of plasma cells by integrin-mediated cell contact signaling (DiLillo et al., 2008) inducing PI3K signaling (Spriel et al., 2012). Here we show that indeed both signaling pathways are essential and complementary for plasma cell survival, and that cell contact-induced PI3K signaling acts through downregulation of FoxO1/3 and blocks activation of caspases 3 and 7, but not 12. In complementation, APRIL signaling blocks activation of caspase 12, but not 3 and 7. Direct contact between the stromal cell and the plasma cell, and APRIL signaling thus in synergy prevent apoptosis of the plasma cell induced by ER and mitochondrial stress.

Analysis of the lifestyle of memory plasma cells has been hampered by their low frequency in the bone marrow, as well as by the fact that they die rapidly, when isolated and cultured *ex vivo* (Cassese et al., 2003), as we confirm here. Cytokines prolonging their survival *ex vivo* have been reported (Cassese et al., 2003; Cocco et al., 2012; Jourdan et al., 2014). Here we describe a culture system mimicking their physiological niche as close as possible. When cultured on ST2 stromal cells, in the presence of the cytokine APRIL, 50 to 80% of *ex vivo* isolated bone marrow plasma cells survived for more than 5 days. The plasma cells maintained their transcriptional identity, with less than 41 transcripts differentially expressed between plasma cells directly isolated from the bone marrow, and those cultured for 3 days.

*In vivo*, it remains unclear, which cells provide the cytokines APRIL and/or BAFF to the plasma cells. Eosinophilic granulocytes have been proposed (Chu et al., 2011), but their role has been questioned (Haberland et al., 2018). We have shown recently, that a subpopulation of bone marrow stromal cells does produce BAFF (Addo et al., 2019). Plasma cells are known to have two receptors for the cytokines APRIL and BAFF, namely TACI and BCMA (O’Connor et al., 2004a; Peperzak, 2013). Among them, BCMA signaling has been suggested to be vital for plasma cells survival in the bone marrow (Benson et al., 2008; O’Connor et al., 2004a). It is known that the BCMA receptor, when activated by APRIL, signals via the TRAF/NF-κB/p38 pathway (Hatzoglou et al., 2000). Similarly, inhibition of the NF-κB pathway blocks survival of plasma cells in the *ex vivo* niche described here. In the presence of stromal cells, APRIL significantly inhibits the activation of caspase 12, but not of caspases 3 and 7. Caspase 12 is localized at the ER and is activated by ER stress, e.g. by an insufficient unfolded protein response (UPR) (Nakagawa et al., 2000). It was previously shown that managing the UPR is essential for plasma cell survival (Iwakoshi, 2003; Pelletier et al., 2006) since they are cells producing several thousand antibody molecules per second (Hibi & Dosch, 1986).

The original notion, that direct cell contact between stromal cell and plasma cell is essential for the survival of plasma cells in the bone marrow, comes from their ablation *in vivo*, by antibodies against integrin α4β1 (VLA-4), a ligand of the stromal cell receptor VCAM1, and integrin αLβ2 (LFA-1), a ligand of the stromal cell receptor ICAM-1, both expressed by bone marrow plasma cells (DiLillo et al., 2008). At a molecular level, it has been demonstrated that integrins can signal via the FAK/PI3K-pathway (Giancotti, 1999). An indication that this may also happen in plasma cells comes from mice deficient for the tetraspanin CD37. In these mice, antibody secreting cells showed impaired clustering of VLA4, and diminished PI3K-signaling *ex vivo* (Spriel et al., 2012). Here we demonstrate by transwell separation of plasma cells and stromal cells in the *ex vivo* niche, that direct contact of bone marrow plasma cells to ST2 stromal cells is required for survival of the plasma cells. The contact induces PI3K signaling in the plasma cells, which is required for their survival. Inhibitors specific for the α-, β-, γ- or δ-subunits of PI3K (p110-α: BYL719, p110-β: GSK2636771, p110-δ: IC-87114, p110-γ: CZC24832), in any combination of three blocked plasma cell survival, suggesting that plasma cells can use any two of the four different PI3K-subunits for survival signaling, in a redundant fashion. Also the PI3K inhibitors LY29400 and Wortmannin block survival of the plasma cells *ex vivo* efficiently. Wortmannin also ablates plasma cells from bone marrow, *in vivo*, with the same efficiency as in the *ex vivo* niche. Considering the different spectra of off-target effects of all those inhibitors, the results strongly suggest that indeed stromal cell-contact induced PI3K signaling is vital for memory plasma cell survival. The cell-contact in part is mediated by ß1-integrin, since siRNA-mediated knock down of ß1 integrin in plasma cells impairs survival of these cells in the *ex vivo* niche, as we show here.

PI3K signaling via activation of AKT, has several downstream targets. One of them is the mechanistic target of rapamycin complex 1 (mTORC1). This has previously been shown to be not involved in memory plasma cell survival (Jones et al., 2016). Others are the transcription factors FoxO1/3. Activated AKT phosphorylates FoxO1/3, thus inactivating them as transcription factors. Inactivation of FoxO1/3 has been shown to be essential for survival and proliferation of activated lymphocytes, although the exact molecular mechanisms remain enigmatic (Haftmann et al., 2012; Stittrich et al., 2010). Here, we show, that inactivation of FoxO1/3 is essential for survival of memory plasma cells, since siRNA-mediated knock down of FoxO1/3 in the plasma cells nearly fully compensates for stromal cell contact in the *ex vivo* niche.

Unlike APRIL, stromal cells block activation of the caspases 3 and 7, but not 12 in bone marrow plasma cells, in the *ex vivo* niche. Caspases 3 and 7 have been reported to be activated upon mitochondrial stress (Lakhani et al., 2006). Inhibition of caspases 3 and 7 by stromal cell contact, and of caspase 12 by APRIL seems to be necessary and sufficient to prevent apoptosis of memory plasma cells in the *ex vivo* niche. Pan-caspase inhibition can compensate for either one, ST2 cells or APRIL. While stromal cell contact blocks the effector caspases of mitochondrial stress induced apoptosis, APRIL blocks the initiator caspase of ER stress induced apoptosis, caspase 12, apparently the two major stress factors limiting the lifetime of memory plasma cells. Also, the expression of the essential survival factor of plasma cells IRF4 (Tellier et al., 2016) is upregulated by stromal cell contact and APRIL in synergy, as we show here. Taken together, we demonstrate that stromal cells are not just organizing memory plasma cell survival niches in the bone marrow. By integrin-mediated cell contact, they actively provide an essential survival signal to memory plasma cells, complementing the survival signal provided by APRIL/BAFF. Both signals efficiently prevent mitochondrial and ER-stress induced apoptosis of memory plasma cells.

## STAR METHODS

- Contact for reagent and resource sharing
- Experimental model
  ○ Mice
- Method details
  ○ Immunization
  ○ *In vivo* treatment with PI3K-inhibitor
  ○ Magnetic isolation of long-lived plasma cells from the bone marrow
  ○ Cell culture of long-lived plasma cells and treatment with inhibitors
  ○ siRNA treatment of plasma cells *in vitro*
  ○ CaspGlow staining
  ○ Flow cytometric measurement of surface and intracellular antigens
  ○ ELISA
  ○ Transwell-Assay
  ○ Gene Expression Microarray Analysis
- Quantification and statistical analysis
- Key resources table

## CONTACT FOR REAGENT AND RESOURCE SHARING

Further information and requests for resources and reagents should be directed to and will be fulfilled by the Lead Contact, Andreas Radbruch.

## EXPERIMENTAL MODELS

### Mice

C57BL/6J wild type strain was purchased from Charles River Laboratories. Mice expressing GFP under the control of the Prdm1 promoter (Blimp-1:GFP) were bred and maintained at the “Bundesinstitut für Risikobewertung” (BfR, Berlin, Germany), a gift from S. Nutt (Walter and Eliza Hall Institute, Melbourne, Australia). All mice were maintained under specific pathogen free conditions. All experiments were performed according to German law for animal protection and with the permission from the local veterinary offices, and in compliance with the guidelines of the Institutional Animal Care and Use Committee.

## METHOD DETAILS

### Immunization

Mice were primed with 100μg 4-hydroxy-3-nitropheylacetyl hapten coupled chicken gamma globulin (NP-CGG) in incomplete Freud’s Adjuvants (IFA) intraperitoneally (i.p.). In total 200μl (100μl of NP-CGG diluted in PBS + 100μl IFA) have been injected per mouse. Mice were challenged twice after the prime with the same injection in cycles of 21 days.

### *In vivo* treatment with PI3K-inhibitor

Immunized mice were used and Wortmannin in DMSO or PBS in DMSO injected i.p. with a total volume of 100μl on day 90, 92 and 94. Concentration of Wortmannin was 1.2mg/kg. Mice were killed by cervical dislocation and analyzed on day 95.

### Magnetic isolation of long-lived plasma cells from the bone marrow

Plasma cells were magnetically isolated from immunized mice more than 30 days after 2^nd^ boost using a two-step protocol, including depletion of B220 and CD49b expressing cells and subsequent positive enrichment of CD138^high^ plasma cells.

### Cell culture of long-lived plasma cells and treatment with inhibitors

Isolated long-lived plasma cells from the bone marrow were cultured in RPMI1640 medium supplemented with 10% FCS, 100 U/ml Penicillin, 100 μg/ml streptomycin, 0.1% β-Mercaptoethanol, 25 mM HEPES buffer and 50 ng/ml multimeric APRIL. Cultures were kept under physiological oxygen levels in a hypoxia chamber with 4.2% O_2_ and 5% CO_2_ at 37°C. Cells were pre-treated with pan PI3K inhibitors at different concentrations including Wortmannin, Ly294002 and to block NF-κB pathway the inhibitor, IKK16, was used.

### siRNA treatment of plasma cells *in vitro*

Isolated long-lived plasma cells from the bone marrow were cultured in Accell Medium with 2 μM siRNA or scr control (Bardua et al., 2018; Haftmann et al., 2015, 2015) and 100 ng/ml multimeric APRIL. After 1 hour RMI1640 medium supplemented with 5% FCS, 200 U/ml Penicillin, 200 μg/ml streptomycin0.2% β-Mercaptoethanol, 50 mM HEPES buffer was added to the cells. Cultures were kept under physiological oxygen levels in a hypoxia chamber with 4.2% O_2_ and 5% CO_2_ at 37°C.

### CaspGlow staining

CaspGlow assays were performed by staining the cells in cell culture medium with the CaspGlow FITC labeled peptide for 30 minutes at 37°C. Cells were washed two times, stained for CD138 and measured by flow cytometry.

### Flow cytometric measurement of surface and intracellular antigens

Single cell suspension was prepared; cells were pre-incubated with uncoupled anti-FCyRII/III antibody to block unspecific binding, followed by direct addition of antibodies for 15 minutes on ice. For staining of intracellular antigens, cells were fixed with PFA and permeabilized with methanol. To prevent unspecific binding, cells were pre-incubated with blocking buffer containing rabbit or rat serum and subsequently stained with primary antibody for 1 hour and, if necessary, with secondary antibody for 30 minutes. Samples were analyzed using a MacsQuant analyzer and FlowJo software. Cytometric procedures followed the recommendations of the “Guidelines for use of flow cytometry and cell sorting in immunological studies” (Cossarizza et al., 2017).

### ELISA

Enzyme-linked immunosorbent assay was used to detect secreted antibodies in supernatants at different time points of the plasma cell culture. Supernatant was incubated in plates coated with unlabeled IgG, IgM or IgA. Isotype specific antibodies were detected using biotinylated IgG, IgM or IgA and streptavidin-peroxidase.

### Transwell-Assay

Isolated plasma cells were cultured in transwell plates with pore size of 0.5μm. Plasma cells were plated at the bottom and stromal cells on top of the transwell culture. As control of direct contact effects plasma cells and stromal cells were both plated on the top of the filter.

### Processing and analysis of oligonucleotide microarray data

Memory plasma cells were isolated by magnetic cell sorting or after 3 days of culture in the *in vitro* system were processed for RNA preparation. RNA was prepared using the RNeasy Micro KIT and hybridized to mouse 430 2 GeneChips according to the whole-transcriptome pico KIT. Raw signals were processed by the affy R package using RMA for normalization (Gautier, Cope, Bolstad, & Irizarry, 2004). Sample similarity was evaluated by Pearson correlation and Principle component analysis based on un-scaled log2 expression values. For the differentially expressed gene analysis the limma R package was used (Ritchie et al., 2015). Genes with an adjusted P-value < 0.05 were considered to be statistically differentially expressed. Microarray data is available through Gene Expression Omnibus (GEO:GSE107206; code for reviewer: sfctqoakblqhpyp).

## QUANTIFICATION AND STATISTICAL ANALYSIS

Data are presented as median. Data was tested for normality and significant differences between two or more groups were determined by performing Mann-Whitney test, Kruskal-Wallis test or ordinary one-way ANOVA. Differences were considered statistically significant when p < 0.05.

## KEY RESOURCES TABLE

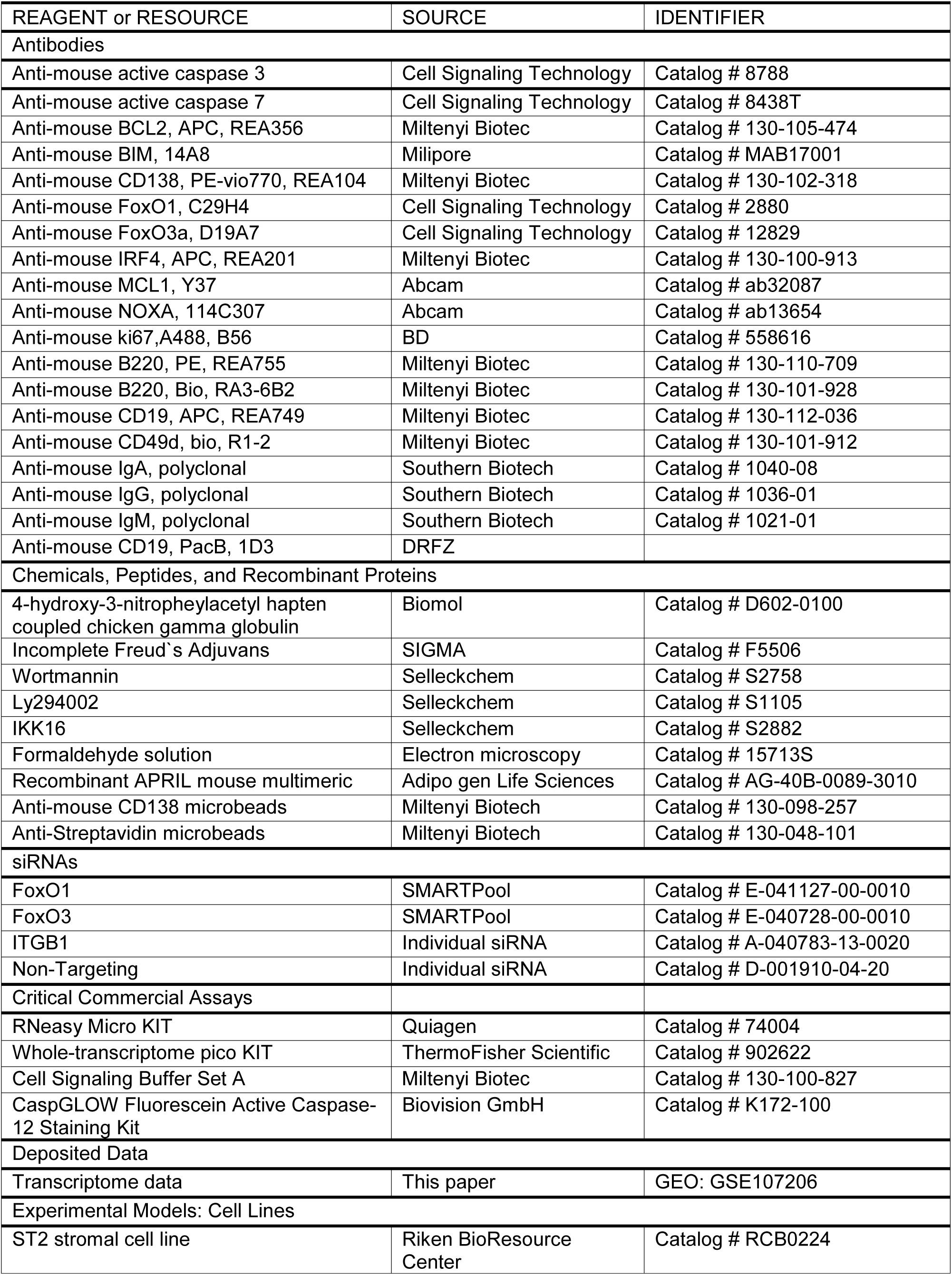
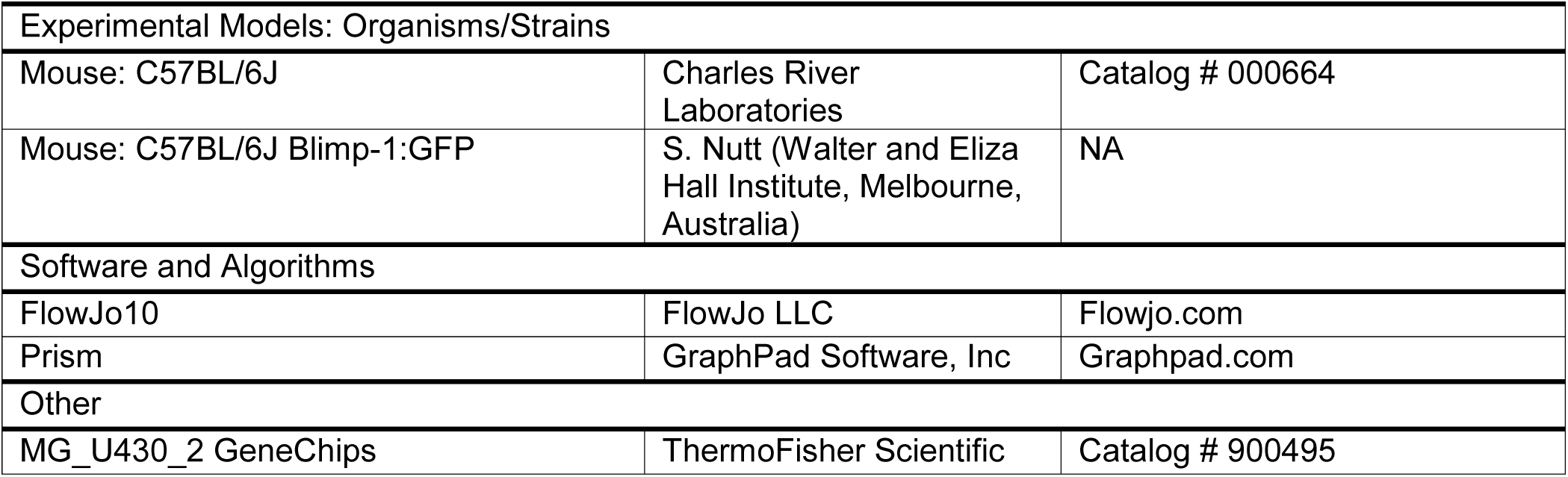

| REAGENT or RESOURCE | SOURCE | IDENTIFIER |
| --- | --- | --- |
| **Antibodies** |  |  |
| Anti-mouse active caspase 3 | Cell Signaling Technology | Catalog # 8788 |
| Anti-mouse active caspase 7 | Cell Signaling Technology | Catalog # 8438T |
| Anti-mouse BCL2, APC, REA356 | Miltenyi Biotec | Catalog # 130-105-474 |
| Anti-mouse BIM, 14A8 | Milipore | Catalog # MAB17001 |
| Anti-mouse CD138, PE-vio770, REA104 | Miltenyi Biotec | Catalog # 130-102-318 |
| Anti-mouse FoxO1, C29H4 | Cell Signaling Technology | Catalog # 2880 |
| Anti-mouse FoxO3a, D19A7 | Cell Signaling Technology | Catalog # 12829 |
| Anti-mouse IRF4, APC, REA201 | Miltenyi Biotec | Catalog # 130-100-913 |
| Anti-mouse MCL1, Y37 | Abcam | Catalog # ab32087 |
| Anti-mouse NOXA, 114C307 | Abcam | Catalog # ab13654 |
| Anti-mouse ki67, A488, B56 | BD | Catalog # 558616 |
| Anti-mouse B220, PE, REA755 | Miltenyi Biotec | Catalog # 130-110-709 |
| Anti-mouse B220, Bio, RA3-6B2 | Miltenyi Biotec | Catalog # 130-101-928 |
| Anti-mouse CD19, APC, REA749 | Miltenyi Biotec | Catalog # 130-112-036 |
| Anti-mouse CD49d, bio, R1-2 | Miltenyi Biotec | Catalog # 130-101-912 |
| Anti-mouse IgA, polyclonal | Southern Biotech | Catalog # 1040-08 |
| Anti-mouse IgG, polyclonal | Southern Biotech | Catalog # 1036-01 |
| Anti-mouse IgM, polyclonal | Southern Biotech | Catalog # 1021-01 |
| Anti-mouse CD19, PacB, 1D3 | DRFZ |  |
| **Chemicals, Peptides, and Recombinant Proteins** |  |  |
| 4-hydroxy-3-nitrophenylacetyl hapten coupled chicken gamma globulin | Biomol | Catalog # D602-0100 |
| Incomplete Freud's Adjuvans | SIGMA | Catalog # F5506 |
| Wortmannin | Selleckchem | Catalog # S2758 |
| Ly294002 | Selleckchem | Catalog # S1105 |
| IKK16 | Selleckchem | Catalog # S2882 |
| Formaldehyde solution | Electron microscopy | Catalog # 15713S |
| Recombinant APRIL mouse multimeric | Adipo gen Life Sciences | Catalog # AG-40B-0089-3010 |
| Anti-mouse CD138 microbeads | Miltenyi Biotec | Catalog # 130-098-257 |
| Anti-Streptavidin microbeads | Miltenyi Biotec | Catalog # 130-048-101 |
| **siRNAs** |  |  |
| FoxO1 | SMARTPool | Catalog # E-041127-00-0010 |
| FoxO3 | SMARTPool | Catalog # E-040728-00-0010 |
| ITGB1 | Individual siRNA | Catalog # A-040783-13-0020 |
| Non-Targeting | Individual siRNA | Catalog # D-001910-04-20 |
| **Critical Commercial Assays** |  |  |
| RNeasy Micro KIT | Qiagen | Catalog # 74004 |
| Whole-transcriptome pico KIT | ThermoFisher Scientific | Catalog # 902622 |
| Cell Signaling Buffer Set A | Miltenyi Biotec | Catalog # 130-100-827 |
| CaspGLOW Fluorescein Active Caspase-12 Staining Kit | Biovision GmbH | Catalog # K172-100 |
| **Deposited Data** |  |  |
| Transcriptome data | This paper | GEO: GSE107206 |
| **Experimental Models: Cell Lines** |  |  |
| ST2 stromal cell line | Riken BioResource Center | Catalog # RCB0224 |
| Experimental Models: Organisms/Strains |  |  |
| Mouse: C57BL/6J | Charles River Laboratories | Catalog # 000664 |
| Mouse: C57BL/6J Blimp-1:GFP | S. Nutt (Walter and Eliza Hall Institute, Melbourne, Australia) | NA |
| Software and Algorithms |  |  |
| FlowJo10 | FlowJo LLC | Flowjo.com |
| Prism | GraphPad Software, Inc | Graphpad.com |
| Other |  |  |
| MG_U430_2 GeneChips | ThermoFisher Scientific | Catalog # 900495 |

## Author contributions

R.C and S.H. designed and performed the experiments, analyzed and interpreted the data and wrote the manuscript. D.M, M.T., C.K., L.H., F.S. carried out and analyzed certain experiments, B.H and F.H. provided mouse strains and helped to write the manuscript, G.P., A.T., P.D., E.M., K.T., F.M., and M.F.M. analyzed data and provided scientific suggestions. H.D.C. and A.R. conceived the study and wrote the manuscript.

## Acknowledgements

Tuula Geske, Heidi Hecker-Kia and Heidi Schliemann we want to thank for expert technical help, Toralf Kaiser and Jenny Kirsch as operators of the flow cytometry core facility (FCCF), Patrick Thiemann and Manuela Ohde for assistance with animal care. This work was supported by European Research Council Advanced Grant IMMEMO (ERC-2010-AdG.20100317 Grant 268987; to A.R.) and by the Deutsche Forschungsgemeinschaft (TRR130; to A.R. and H.D.C.), Innovative Medicines Initiative 2 Joint Undertaking under grant agreement no. 777357. G.P. and M.F.M were supported by the state of Berlin and the “European Regional Development Fund” (ERDF 2014–2020, EFRE 1.8/11, Deutsches Rheuma-Forschungszentrum Berlin). F.S. was supported by Osteoimmune, a FP7 Marie Curie Initial Training Network (FP7-PEOPLE-2011-ITN-289150). E.M. was supported by the Deutsche Forschungsgemeinschaft (MO 2934/1-1-1). This work was supported by the Leibniz ScienceCampus Chronic Inflammation (www.chronische-entzuendung.org).

