## Supplemental Figures for "Stromal cell-contact dependent PI3K and APRIL induced NF-κB signaling complement each other to prevent mitochondrial- and endoplasmic reticulum stress induced cell death of bone marrow plasma cells"

S1.

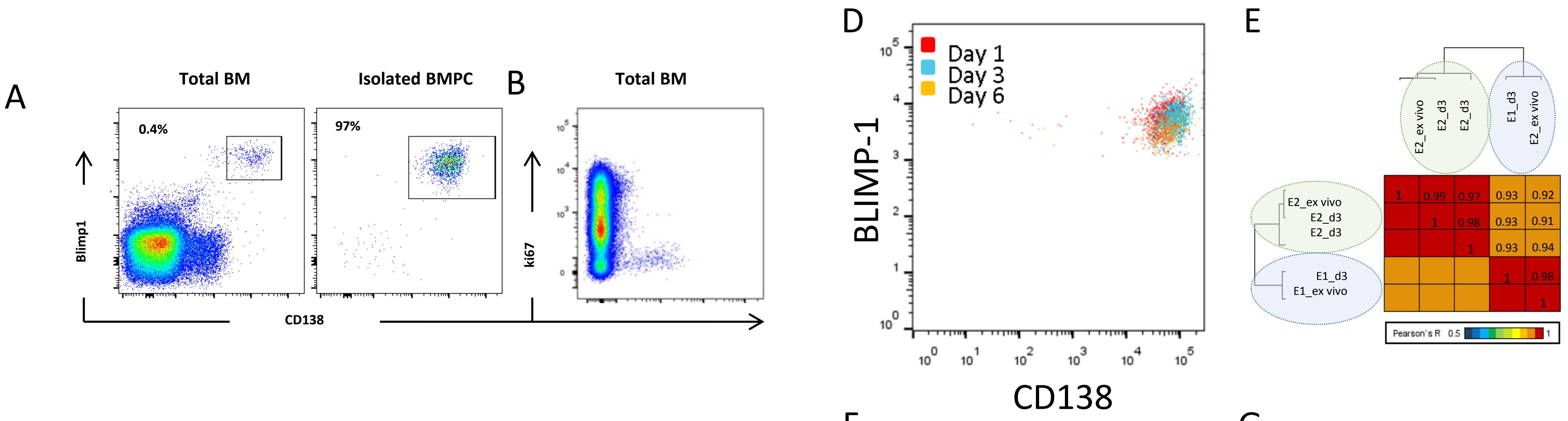

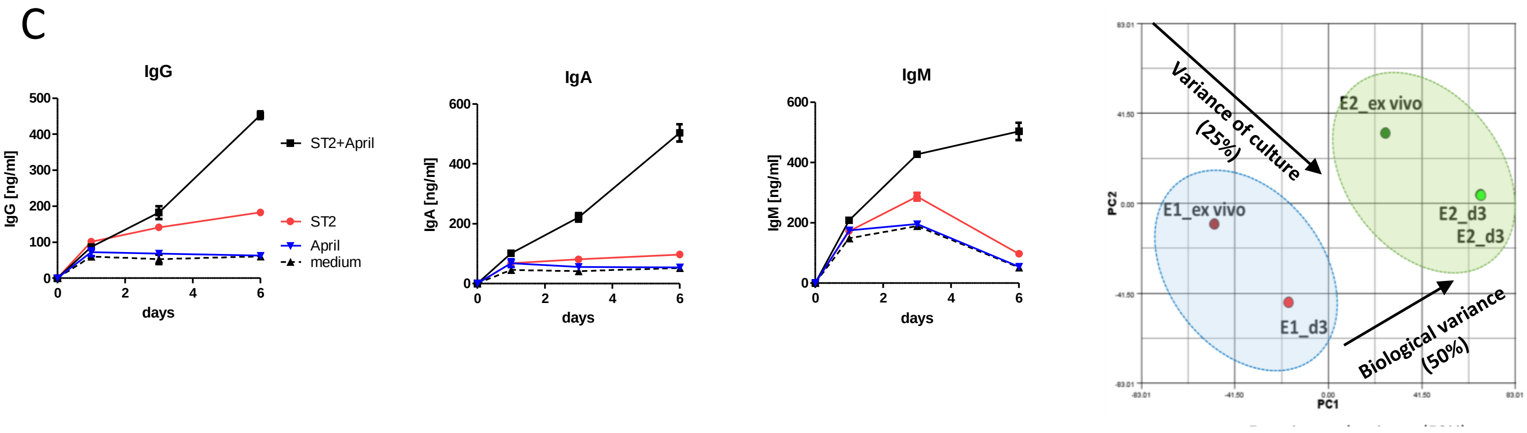

G

| Symbol | mean ex vivo | mean d3 | FC |
|---|---|---|---|
| Prdm1/Blimp-1 | 3864.665 | 3035.403 | 0.78 |
| Sdc1/CD138 | 3339.13 | 3278.153 | 0.98 |
| Bcl2 | 1347.885 | 1027.57 | 0.76 |
| Bcl2l11/Bim | 1051.34 | 1206.74 | 1.14 |
| Pmaip1/Noxa | 333.795 | 371.78 | 1.11 |
| Mcl1 | 1928.82 | 2171.1 | 1.12 |
| Irf4 | 5933.32 | 6511.95 | 1.09 |
| Foxo1 | 731.06 | 937.07 | 1.28 |
| Foxo3 | 2421.92 | 2544.27 | 1.05 |

**S1. Bone marrow memory plasma cells maintain their transcriptional profile and remain functional during *in vitro* culture.**
(A) Original fraction of bone marrow from immunized C57BL/6J mice containing 0.4% of CD138$^{++}$ plasma cells and purity of isolated plasma cells co-expressing BLIMP-1. (B) Ki-67 versus CD138 staining of total bone marrow. (C) Quantification of IgG, IgA and IgM in the supernatant of cultured plasma cells taken at the indicated time points by ELISA (n=1). (D) Expression of CD138 and BLIMP-1 in plasma cells cultured with ST2 in the presence of APRIL and measured on day 1, 3 and 6 of culture. (E) Pearson correlation of the global transcriptome of *ex vivo* isolated and 3 days cultured plasma cells. (F) Principal component analysis of two individual gene expression analyses of plasma cells ex vivo isolated and after 3 day culture. (G) Mean expression and fold-change of plasma cell specific genes and survival genes in *ex vivo* and 3 days cultured plasma cells.

S2.

### S2. Differentially expressed genes of *ex vivo* and cultured plasma cells.

Transcriptomes of memory plasma cells *ex vivo* and after 3 days of culture were analyzed for differentially expressed genes. The table shows statistically differentially expressed genes with adjusted p-value < 0.05 and the corresponding mean expression values.

| | Affy_ID | Symbol | UP:Protein_Name | Limma_q | Mean_Ex vivo plasma cells | Mean_cultured plasma cells | fold-change |
|---|---|---|---|---|---|---|---|
| Group A | 1423100_at | Fos | Proto-oncogene c-Fos; | 1.49E-03 | 8,917.44 | *not present* | |
| | 1422134_at | Fosb | Protein fosB; | 1.49E-03 | 4,600.63 | *not present* | |
| | 1417394_at | Klf4 | Krueppel-like factor 4; | 3.71E-02 | 372.06 | *not present* | |
| | 1418243_at | Fcna | Ficolin-1; | 3.75E-02 | 252.85 | *not present* | |
| | 1420166_at | Dntt | DNA nucleotidylexotransferase; | 4.71E-02 | 144.31 | *not present* | |
| | 1417063_at | C1qb | Complement C1q subcomponent subunit B; | 3.71E-02 | 129.39 | *not present* | |
| | 1417409_at | Jun | Transcription factor AP-1; | 3.71E-02 | 4,521.24 | 185.9 | -24 |
| | 1448830_at | Dusp1 | Dual specificity protein phosphatase 1; | 3.26E-03 | 2,884.55 | 140.41 | -21 |
| | 1448694_at | Jun | Transcription factor AP-1; | 1.46E-02 | 2,537.95 | 205.33 | -12 |
| | 1457404_at | Nfkbiz | NF-kappa-B inhibitor zeta; | 3.84E-02 | 580.27 | 116.43 | -5 |
| | 1430979_a_at | Prdx2 | Peroxiredoxin-2; | 4.71E-02 | 1,103.37 | 355.27 | -3 |
| | 1424854_at | Hist1h4i | Histone H4; | 3.75E-02 | 295.92 | 115.42 | -3 |
| | 1422632_at | Ctsw | Cathepsin W; | 4.71E-02 | 982.67 | 400.7 | -2 |
| Group B | 1415918_a_at | Tpi1 | Triosephosphate isomerase; | 3.71E-02 | 400.97 | 3,633.08 | 9 |
| | 1452927_x_at | Tpi1 | Triosephosphate isomerase; | 3.71E-02 | 518.91 | 3,964.05 | 8 |
| | 1419030_at | Ero1l | ERO1-like protein alpha; | 3.80E-02 | 372.59 | 1,884.26 | 5 |
| | 1433930_at | Hpse | Heparanase; | 3.71E-02 | 231.81 | 1,102.99 | 5 |
| | 1451461_a_at | Aldoc | Fructose-bisphosphate aldolase C; | 3.71E-02 | 276.27 | 1,212.16 | 4 |
| | 1422470_at | Bnip3 | BCL2/adenovirus E1B 19 kDa protein-interacting protein 3; | 3.71E-02 | 224.54 | 954.42 | 4 |
| | 1439148_a_at | Pfkl | ATP-dependent 6-phosphofructokinase, liver type ; | 3.75E-02 | 390.51 | 1,579.83 | 4 |
| | 1449324_at | Ero1l | ERO1-like protein alpha; | 4.71E-02 | 651.93 | 2,260.04 | 3 |
| | 1421057_at | Dnase1l3 | Deoxyribonuclease gamma; | 3.71E-02 | 183.88 | 616.78 | 3 |
| | 1452094_at | P4ha1 | Prolyl 4-hydroxylase subunit alpha-1; | 4.71E-02 | 361.28 | 1,168.42 | 3 |
| | 1451895_a_at | Dusp9 | Dual specificity protein phosphatase ; | 3.71E-02 | 319.3 | 951.66 | 3 |
| | 1417864_at | Pgk1 | Phosphoglycerate kinase 1; | 4.71E-02 | 2,606.08 | 7,562.93 | 3 |
| | 1433604_x_at | Aldoa | Fructose-bisphosphate aldolase; | 3.71E-02 | 2,974.82 | 8,013.74 | 3 |
| | 1418129_at | Dhcr24 | Delta(24)-sterol reductase; | 4.71E-02 | 332.98 | 837.46 | 3 |
| | 1429781_s_at | Ccdc39 | Coiled-coil domain-containing protein 39; | 4.30E-02 | 247.12 | 586.02 | 2 |
| Group C | 1448823_at | Cxcl12 | Stromal cell-derived factor 1; | 4.26E-02 | *not present* | 808.34 | |
| | 1438945_x_at | Gja1 | Gap junction alpha-1 protein; | 3.71E-02 | *not present* | 453.34 | |
| | 1448433_a_at | Pcolce | Procollagen C-endopeptidase enhancer 1; | 3.71E-02 | *not present* | 383.29 | |
| | 1452250_a_at | Col6a2 | Collagen alpha-2(VI) chain; | 3.71E-02 | *not present* | 342.52 | |
| | 1424770_at | Cald1 | H-caldesmon ; | 4.26E-02 | *not present* | 272.95 | |
| | 1434679_at | Ncan | Neurocan core protein; | 4.71E-02 | *not present* | 246.65 | |
| | 1449145_a_at | Cav1 | Caveolin-1; | 2.14E-02 | *not present* | 130.35 | |
| | 1448494_at | Gas1 | Growth arrest-specific protein 1; | 4.12E-02 | *not present* | 121.73 | |
| | 1425028_a_at | Tpm2 | Tropomyosin beta chain; | 3.75E-02 | *not present* | 120.12 | |
| | 1416686_at | Plod2 | Procollagen-lysine,2-oxoglutarate 5-dioxygenase 2; | 4.71E-02 | *not present* | 101.73 | |
| | 1424114_s_at | Lamb1 | Laminin subunit beta-1; | 3.80E-02 | *not present* | 71.56 | |
| | 1429479_at | Magef1 | Putative uncharacterized protein ; | 4.12E-02 | *not present* | 51.38 | |

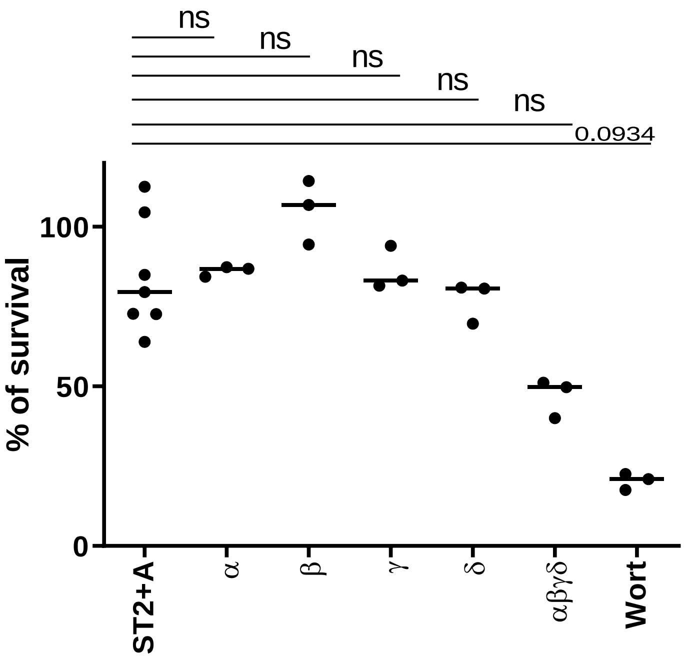

**S3: Subunit specific inhibition of PI3K.**
Survival of memory plasma cells treated with inhibitors against different subunits of PI3K or Wortmannin as control. Cells were enumerated by flow cytometry on day 1 of culture.

S5:

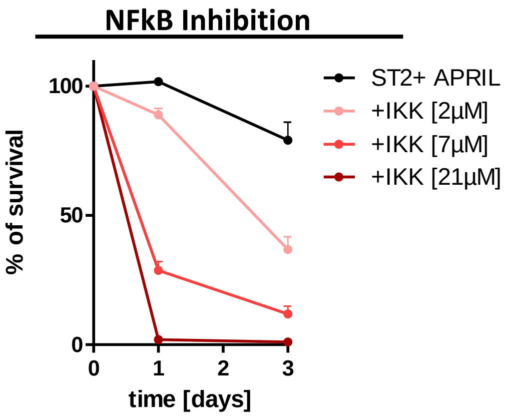

**S4. Memory plasma cell survival depends on NF-kB signaling.**
Survival of memory plasma cells pre-treated with the irreversible inhibitor IKK-16. Cells were enumerated by flow cytometry on day 1 and 3 of culture.
